## Supplementary material for "Brief oxygen exposure after traumatic brain injury speeds recovery and promotes adaptive chronic endoplasmic reticulum stress responses": Sup. Fig.

### 1 Supplemental Findings

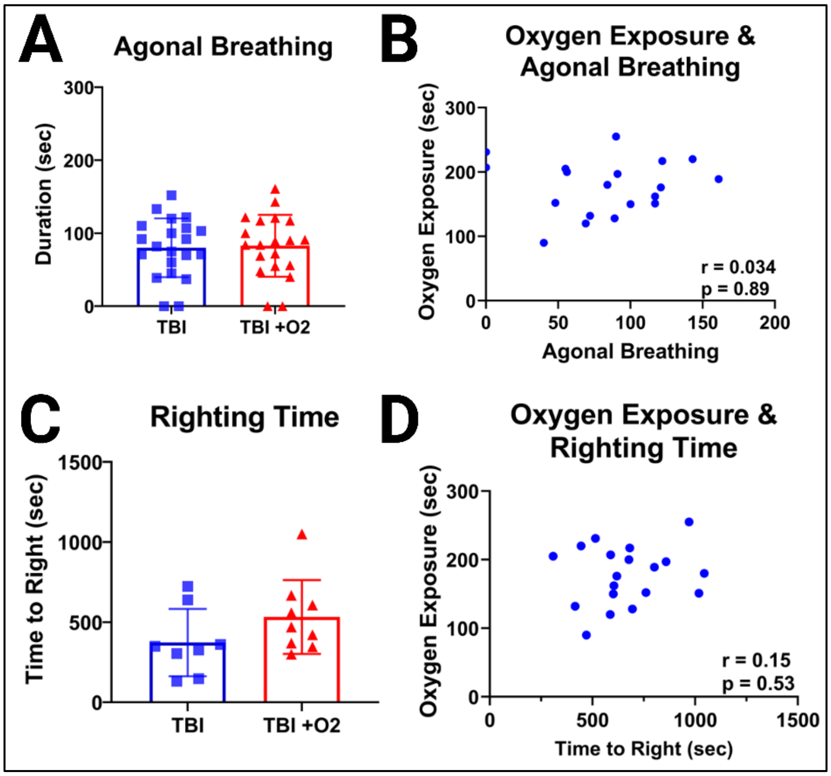


***Supplementary Figure 1. Breathing and oxygen statistics.*** *Agonal breathing (sec) was no different between mice that recovered in room air or in the oxygen chamber, and (B) there was no correlation between time spent in the chamber and cessation of agonal breathing. (C) Righting time was also no different between injury groups, (D) and oxygen exposure duration did not correlate with time to right.*

#### 1.1 Oxygen exposure does not prevent degeneration but may accelerate recovery, supported through similar patterns in projection regions of the optic tract.

In the dorsal lateral geniculate nucleus, a major projection region of the optic nerve, we found a main effect of injury ($F_{2,49}$=26.96, p<0.001) and a significant interaction ($F_{2,49}$=4.15, p=0.02; Sup. Fig. 2a-c). Post-hoc Tuckey's tests showed significant differences between sham and TBI +$O_2$ (p<0.001) and TBI versus TBI +$O_2$ (p<0.001) with mice given oxygen having the highest expression of FJ-C. At day 30, both injured groups were significantly higher than sham (p<0.001). By day 30, TBI mice had significantly reduced FJ expression compared to day 7 (p=0.03) while mice given $O_2$ had similar levels at both times (p=0.9). There was also a main effect of injury ($F_{2,50}$=3.95, p=0.03) in the ventral LGN with injured mice presenting with increased FJ staining at both time points regardless of oxygen exposure (Sup. Fig. 2d).

In the superior colliculi there was a main effect of injury ($F_{2,47}$=20.64, p<0.001; Sup. Fig. 2e-g). Pot-hoc tests revealed significant increases in FJ in TBI mice compared to controls at days 7 (p=0.01) and 30 (p<0.001), this increase was significantly reduced in TBI mice by day 30 (p=0.01). Although there was positive staining in $O_2$ mice, it was not significantly higher than sham at 7 (p=0.5) or 30 days (p=0.14), however, levels were significantly lower than TBI mice 30 days post injury (p=0.04).

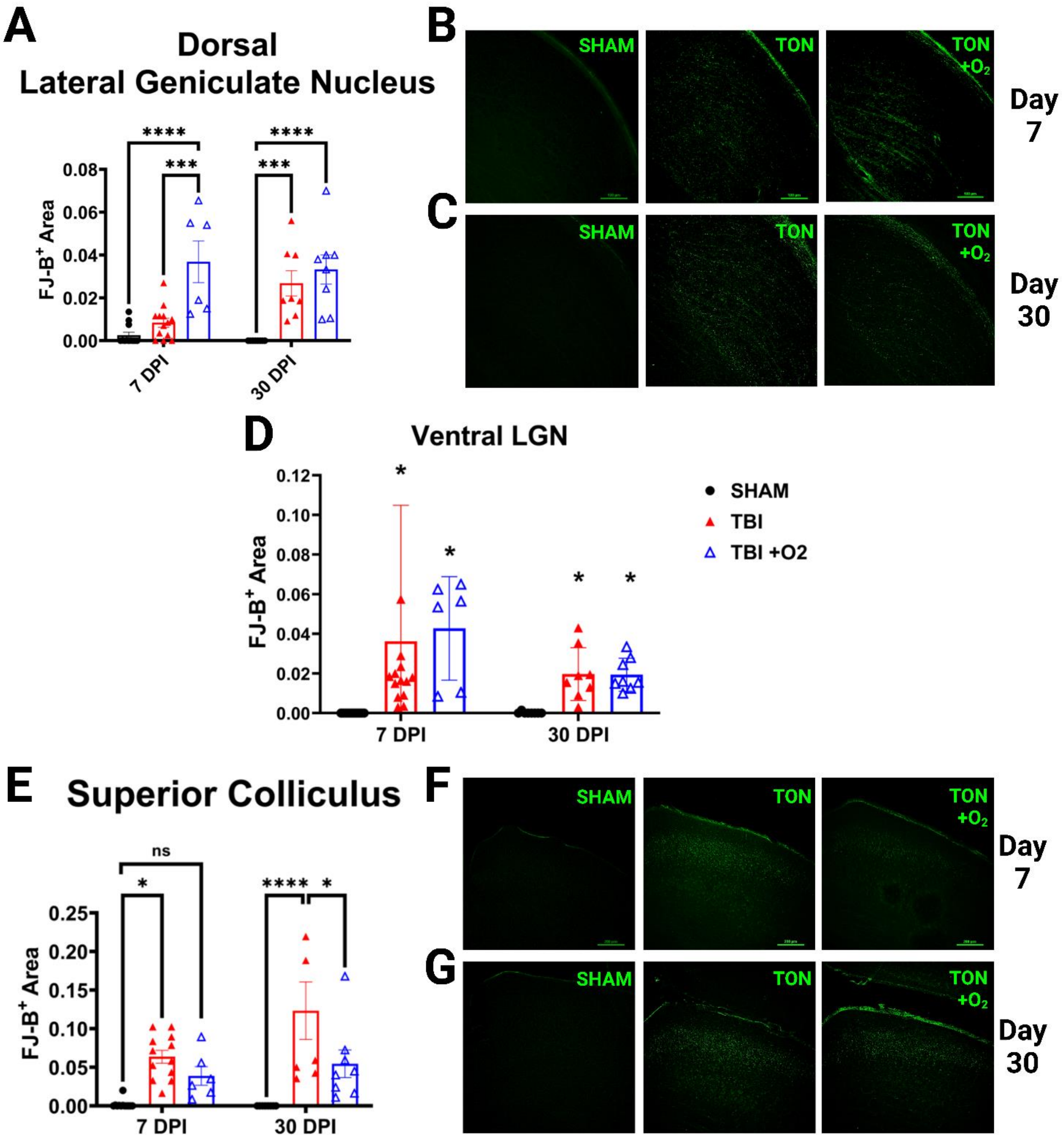


***Supplementary Figure 2. Axonal degeneration in the lateral geniculate nucleus and superior colliculi.*** *(A) the dorsal part of the LGN showed significantly more degeneration in the group given oxygen at 7 DPI, but by day 30- both TBI groups maintained similar levels of FJ-B staining. (B) show representative photomicrographs of the dLGN in 7-day tissue and (C) 30-day tissue. (D) There were no effects of oxygen in the vLGN. (E) In the superior colliculi we, again see potential protection in the oxygen group who, despite having positive staining when there was none in sham mice, were not significantly different from the shams. (F) show representative photomicrographs of the SC in 7-day tissue and (G) 30-day tissue.*

Other regions we examined also showed significant increases in FJ staining in both injured groups at both 7 and 30 days with no effects of time on FJ levels. These regions included arms of the optic tract: the brachium of the superior colliculi (BoSC) and the accessory optic system nuclei (i.e., medial and dorsal terminal nuclei). We also included the nucleus of the optic tract, which is a secondary region of visual input, and the anterior commissure (ACC), which sends some visual information between hemispheres. The anterior commissure was the only of these other regions to reveal an effect of time via a significant interaction ($F_{2,47}$=4.5, p=0.02) such that mice given oxygen have positive FJ staining on day 30, which was both significantly higher than 30-day shams (p=0.01) and TBI +$O_2$ mice on day 7 (p=0.02, Sup. Fig 3e). We have included graphical results in sup. fig. 3. There was no positive staining in the visual cortex or any other cortical region, as we have previously reported in our model.

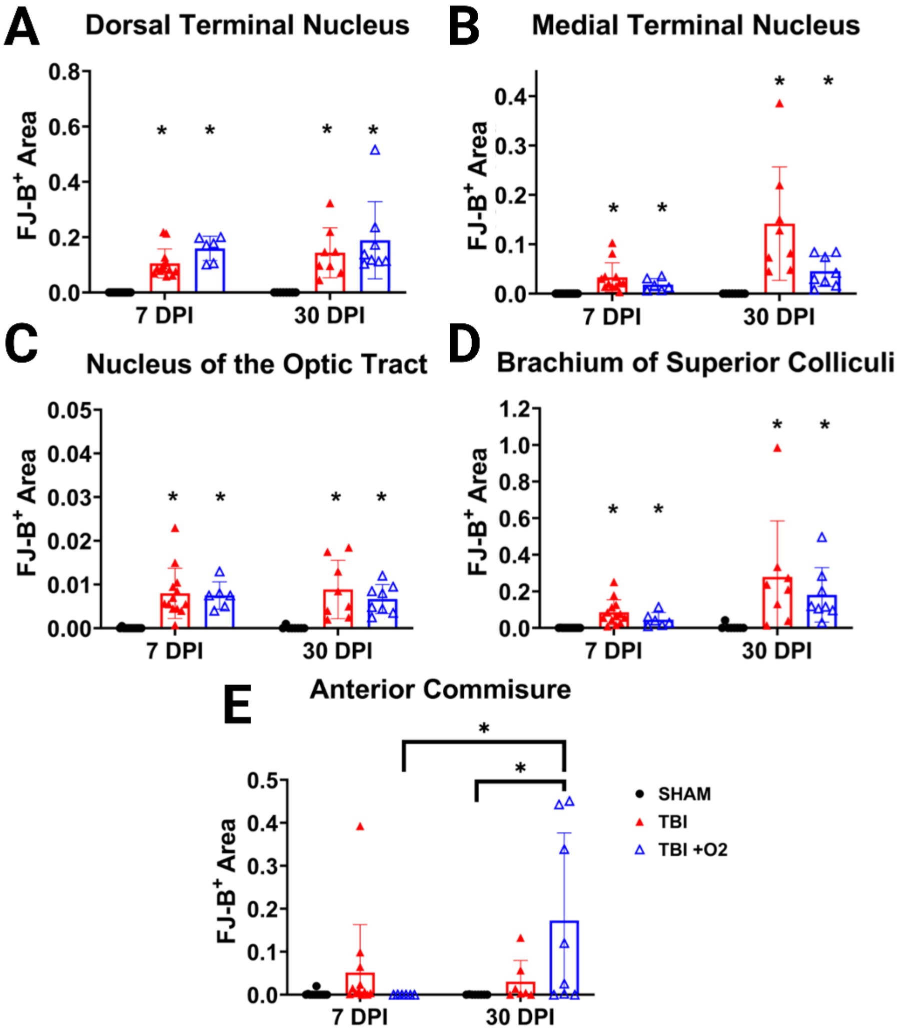


***Supplementary Figure 3. Axonal degeneration in other optic regions is unaffected by oxygen except for the anterior commissure.*** *FJ-B positive staining was significantly present in the (A) dorsal and (B) terminal nuclei of the accessory optic system as well as in the (C) NOT and (D) BoSC. € In the anterior commissure, a major white matter tract bridging left and right hemispheres, only mice given oxygen had ay positive staining at 30 days post injury*

#### 1.2 Astroglial reactivity is significantly reduced in the brain following $O_2$ exposure.

In the dLGN 7 days post injury all groups were different from each other ($F_{2,32}$=73.8, p<0.001). Although both injured groups had higher MFI than sham mice, TBI +$O_2$ mice had significantly lower MFI than TBI mice. By day 30, both injury groups were still significantly higher than sham ($F_{2,24}$=15.6, p<0.001) but MFI levels were similar between injury groups (p=0.8). Results were similar in the vLGN where all groups were different from each other at day 7 ($F_{2,32}$=51.4, p<0.001) and by day 30 injured mice had significantly higher astroglial MFI than sham ($F_{2,24}$=14.7, p<0.001) but injured groups were no longer different from each other (p=0.5). Likewise, mice given oxygen had significantly lower GFAP MFI than TBI mice alone in the superior colliculi (p=0.005) despite still showing elevated MFI compared to sham ($F_{2,31}$=30.1, p<0.001) at day 7, and this decreased MFI in TBI +$O_2$ mice was no longer present 30 days post injury (p=0.4).

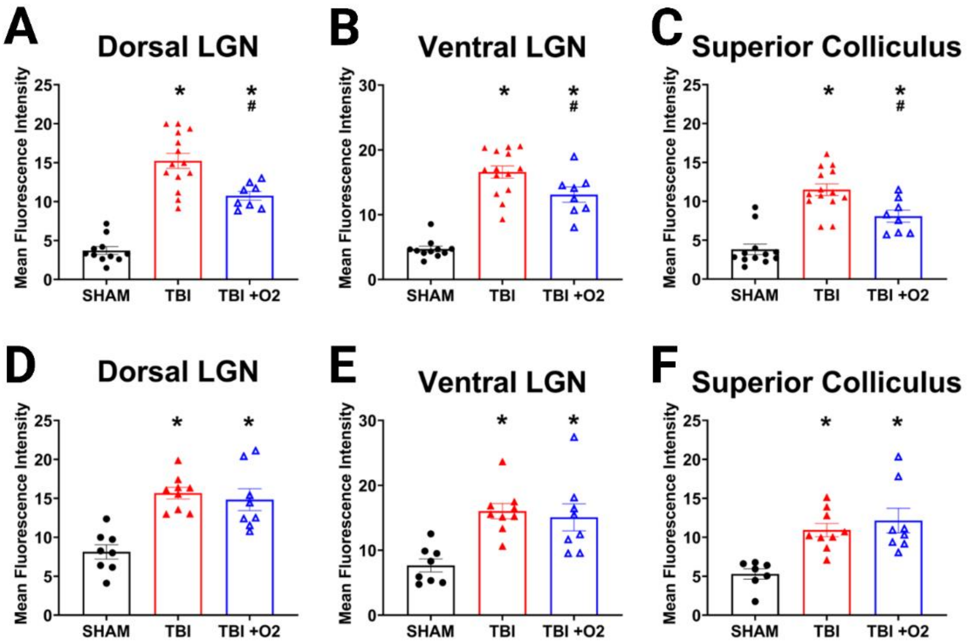


***Supplemental Figure 4. GFAP in optic projection regions.*** *Oxygen significantly reduced GFAP fluorescence intensity in the (A) dorsal and (B) ventral LGN and in the (C) superior colliculus acutely in 7-day tissue, but had no effects at 30 days (D-F).*

Interestingly, we showed significant glial reactivity increased in TBI mice only (p=0.4) in day 30 visual cortex that had not been present in day 7 tissue (p=0.7). We also examined the accessory optic system which is dived into two white matter branches of the optic nerve, the dorsal/lateral and medial terminal nuclei. As with other regions, the dorsal terminal nuclei showed significantly higher MFI in both injured groups ($F_{2,32}$=38.9, p<0.001) 7 DPI. Retinas collected on day 30 showed significantly increased MFI in TBI mice (p=0.02) alone. In the second accessory branch, the medial nuclei, both injured groups higher than sham ($F_{2,32}$=45.1, p<0.001), but this was no longer the case by day 30 (p=0.07).

We found no effects in the suprachiasmatic nucleus at 7 (p=0.5) or 30 days (p=0.5), which is in line with previous studies from our lab (data not shown). In other vision associated areas including the BoSC, nucleus of the optic tract, and ACC we also found significant effects of oxygen. Although there were no acute effects of oxygen in 7-day tissue despite injury effects ($F_{2,31}$=9.5, p<0.001), by day 30 TBI mice had significantly higher MFI than sham (p=0.006), while TBI +$O_2$ mice were no longer different from sham (p=0.2; $F_{2,23}$=6.4, p=0.007). In seven-day tissue, the MFI in the NOT was significantly lower in TBI +$O_2$ mice compared with TBI mice (p=0.02), and mice given oxygen were not different from shams (p=0.2). At 30 days, these effects of oxygen were no longer present and GFAP immunoreactivity remained significantly higher in both injured groups compared to shams ($F_{2,23}$=10.9, p<0.001).

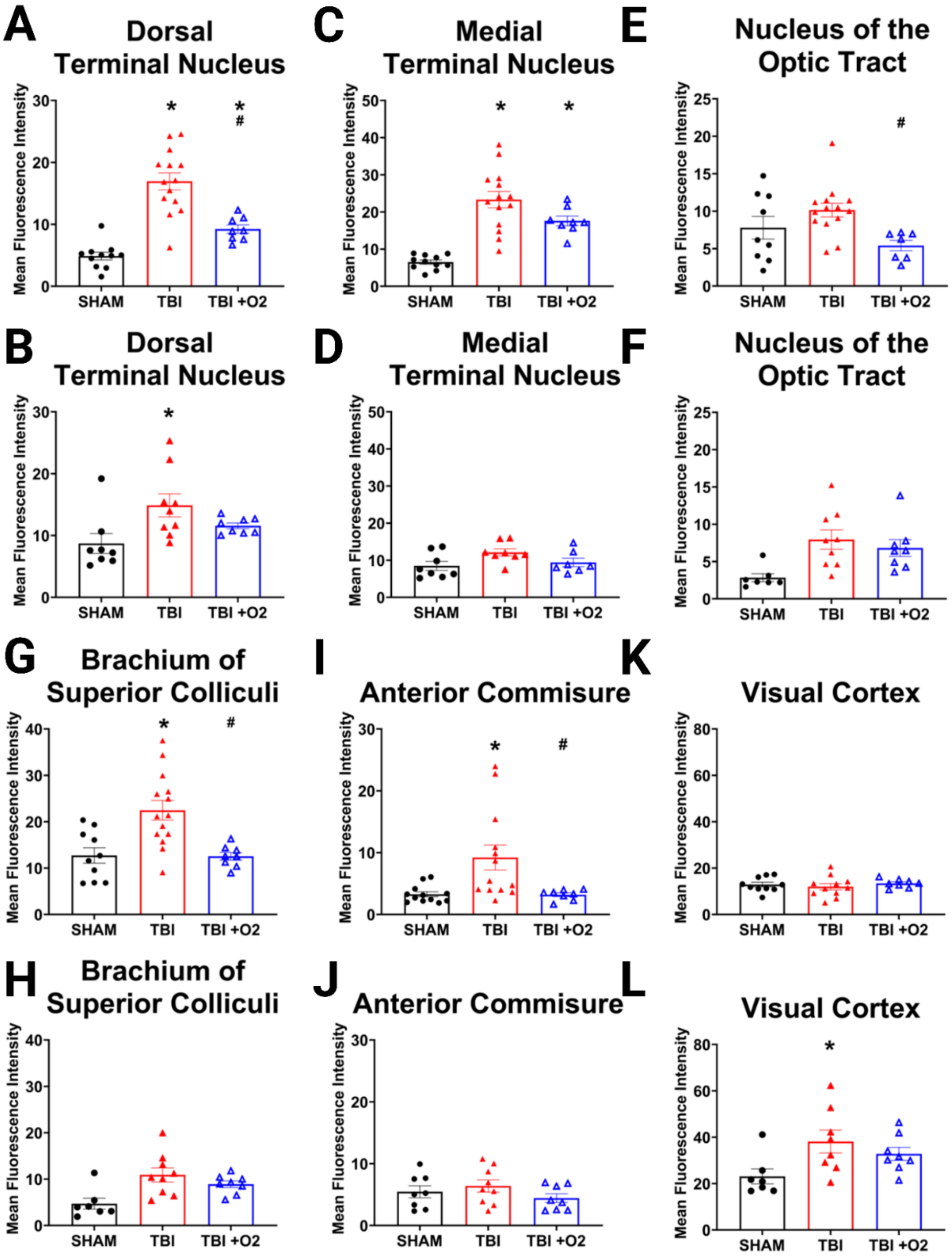


***Supplementary Figure 5. GFAP in other optic regions.*** *In the Dorsal terminal nucleus oxygen significantly reduced GFAP reactivity at 7-days (A) and $O_2$ mice were no longer different from sham at 30 days (B). Astrogliosis was also present in injured mice in the ventral terminal nucleus at 7 days (C) but not at 30 days (D). € TBI +O2 mice at 7 days actually*

*presented with significantly less GFAP MFI than even control mice, but (F) there were no differences in 30-day tissue. (G) oxygen was also beneficial in the BoSC at 7 days with (H) with no differences at 30 days. The same was true of the Anterior Commissure (I-J) with no gliosis present at all in the visual cortex at 7 (K) or 30 (L) days.*

Although data failed equal variance, oxygen prevented a significant change in GFAP MFI in another white matter tract, the ACC (p=1) by day 7. TBI mice had significantly higher MFI than sham (p=0.005) and TBI +$O_2$ mice (p=0.03). There was no longer an increase in MFI in any mice by day 30 (F1,15=0.7, p=0.4).

##### 1.3 The PERK pathway is sub-acutely elevated after TBI, but oxygen reduces long-term expression of apoptotic markers.

Post-hoc probing of retinal tissue for the PERK arm's feedback machinery, growth arrest and DNA damage-inducible protein (GADD34), did not reveal any significant findings ($F_{2,26}$=1.8 p=0.18, Sup. Fig. 6). Of note there were two unidentifiable bands at ~43 and 35 kDa. GADD34 is a relatively uncharacterized protein, but most-importantly, it's primary function is to bind to PP1 in order to dephosphorylate eIF2a.

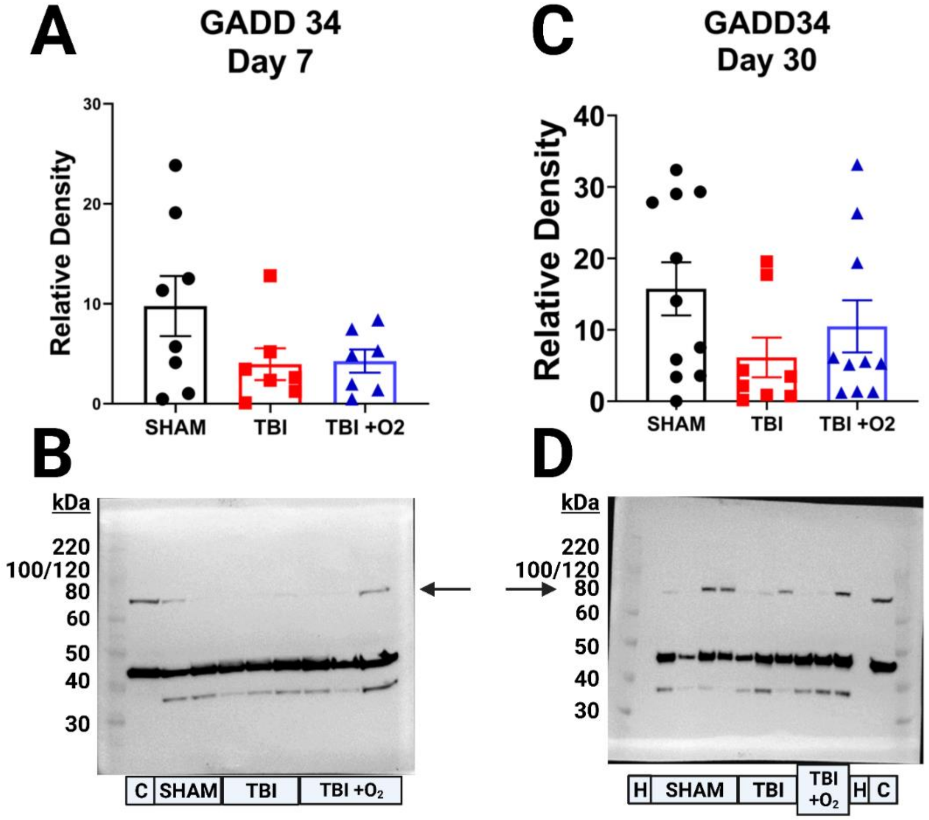


***Supplementary Figure 6. Reduced P-eIF2a is not the results of GADD34.*** *(A) There were no significant changes in GADD34 protein expression in the retina at day 7. (B) Representative western blot of GADD 34 (~78kDa) with two unknown bands.(C) There were also no changes in 30-day tissue. (D) Shows a representative blot.*
